# Behavioural mechanisms underlying the evolution of cooperative burrowing in *Peromyscus* mice

**DOI:** 10.1101/731984

**Authors:** Nicole L. Bedford, Jesse N. Weber, Wenfei Tong, Felix Baier, Ariana Kam, Rebecca A. Greenberg, Hopi E. Hoekstra

## Abstract

While some behaviours are largely fixed and invariant, others can respond flexibly to different social contexts. Here, we leverage the unique burrowing behaviour of deer mice (genus *Peromyscus*) to investigate if and how individuals of three species adapt their behaviour when digging individually versus with partners. First, we find that pairs of mice from monogamous (*P. polionotus*) but not promiscuous (*P. maniculatus, P. leucopus*) species cooperatively construct burrows that are approximately twice as long as those dug by individuals and similar in size to burrows found in the wild. However, the length of burrows built by *P. polionotus* pairs differs: opposite-sex pairs construct longer burrows than same-sex pairs. By designing a novel behavioural assay in which we can observe and measure burrowing behaviour directly, we find that longer burrows are achieved not by changing individual behaviour, but instead because opposite-sex pairs are more socially cohesive and thus more likely to dig simultaneously, which is a more efficient mode of burrow elongation. Thus, across social contexts, individual burrowing behaviour appears largely invariant, even when the resultant burrow from pairs of mice differs from expectation based on individual behaviour, underscoring the fixed nature of burrowing behaviour in *Peromyscus* mice.

## Introduction

Animal behaviours, like other heritable traits, are subject to evolution by natural selection, which can promote either the rigid expression of a given behaviour or, at the other extreme, flexibility in behaviour. For some species, the reliable execution of a fixed response can be advantageous, while for others, the ability to flexibly adapt to changing contexts may be beneficial. Ernst Mayr^1^ described these fixed and flexible behaviours as being governed by “closed” (i.e., impervious to modification) versus “open” genetic programs (i.e., subject to external influences). Interestingly, the same behaviour may be closed in one species, yet open in another. For instance, birdsong in the common cuckoo – a species reared by heterospecific foster parents – is largely invariant^2^; whereas European starlings are open-ended learners and can modify their song well into adulthood^3,4^. Mayr^1^ suggested that open behavioural programs may be more likely to evolve in more social species, in which flexibility in complex social environments may be especially beneficial. However, both comparative studies – in which homologous behaviours are contrasted across species – and studies of individual behaviours across different social contexts are lacking.

Animal architectures – such as termite mounds, bird nests, or rodent burrows – are stereotyped structures that can aid in species identification^5,6^ or the reconstruction of phylogenetic histories^7,8^ and, as such, are prime candidates for comparative studies of homologous behaviour^9,10^. The burrows built by North American deer mice (genus *Peromyscus*) are highly stereotyped and can be readily quantified in the laboratory^11-13^. For example, the oldfield mouse (*P. polionotus*) constructs “complex” burrows consisting of a long entrance tunnel, central nest chamber, and upward-sloping escape tunnel^14,15^. By contrast, its sister species (*P. maniculatus*) and an outgroup (*P. leucopus*) produce relatively simple burrows comprised of only a short entrance tunnel and terminal nest chamber^11,12^. Moreover, in many rodents, burrows are an important site for social interactions^16^, but the type and degree of social interaction can vary greatly across species. In the wild, *P. polionotus* are both socially and genetically monogamous^17-19^, whereas *P. maniculatus* and *P. leucopus* are highly promiscuous^20^. Here, we asked if individual behaviour – burrow construction – changes with different social contexts and if this response varies across species with different mating systems.

Using a combination of novel behavioural assays – capturing both variation in burrow architecture and individual digging behaviour – we tested whether mice of three *Peromyscus* species alter their behaviour when burrowing alone versus with a partner. More specifically, we compared the lengths of burrows constructed by individuals and pairs, and further compared the burrows constructed by same- and opposite-sex pairs. We show that only mice of the monogamous species (*P. polionotus*) dig cooperatively, and that while jointly-constructed burrows are longer than individual burrows, their length differs depending on the sex of one’s digging partner. These differences, however, are not due to changes in individual burrowing behaviours, such as time spent digging, but rather due to more simultaneous digging – a more efficient mode of burrowing.

## Results

### Effect of social context on burrow length

When tested as individuals, *Peromyscus* mice build burrows that are highly stereotyped and species-specific in size and shape^12,21^ (Fig. 1a); however, it remains unknown if burrows are constructed by pairs of mice, and if so, how they may differ in size or shape from individual burrows. To address these questions, we first compared the length of the longest burrow per trial produced by both individuals and pairs of mice in large, sand-filled enclosures (see Methods). While species-specific burrow shape remained unaltered between individual and pair trials, we found significant variation in burrow length. Using linear mixed-effects models (LMMs) that control for sex and mouse identity, we found that individual and pair trials resulted in equivalent burrows for both *P. leucopus* and *P. maniculatus*, while in *P. polionotus*, pairs of mice produced significantly longer burrows than individuals (Fig. 1b, LMMs, *P. leu*: *P* = 0.631; *P. man*: *P* = 0.071; *P. pol*: *P* < 0.001). In *P. polionotus*, pairs dug 82% longer burrows, on average, than individuals (64.3 ± 4.2 cm vs. 35.3 ± 2.1 cm). Thus, the response to social context varies by species: *P. polionotus* produces longer burrows when digging in pairs, whereas its congeners (*P. maniculatus* and *P. leucopus*) do not.

**Figure 1.**
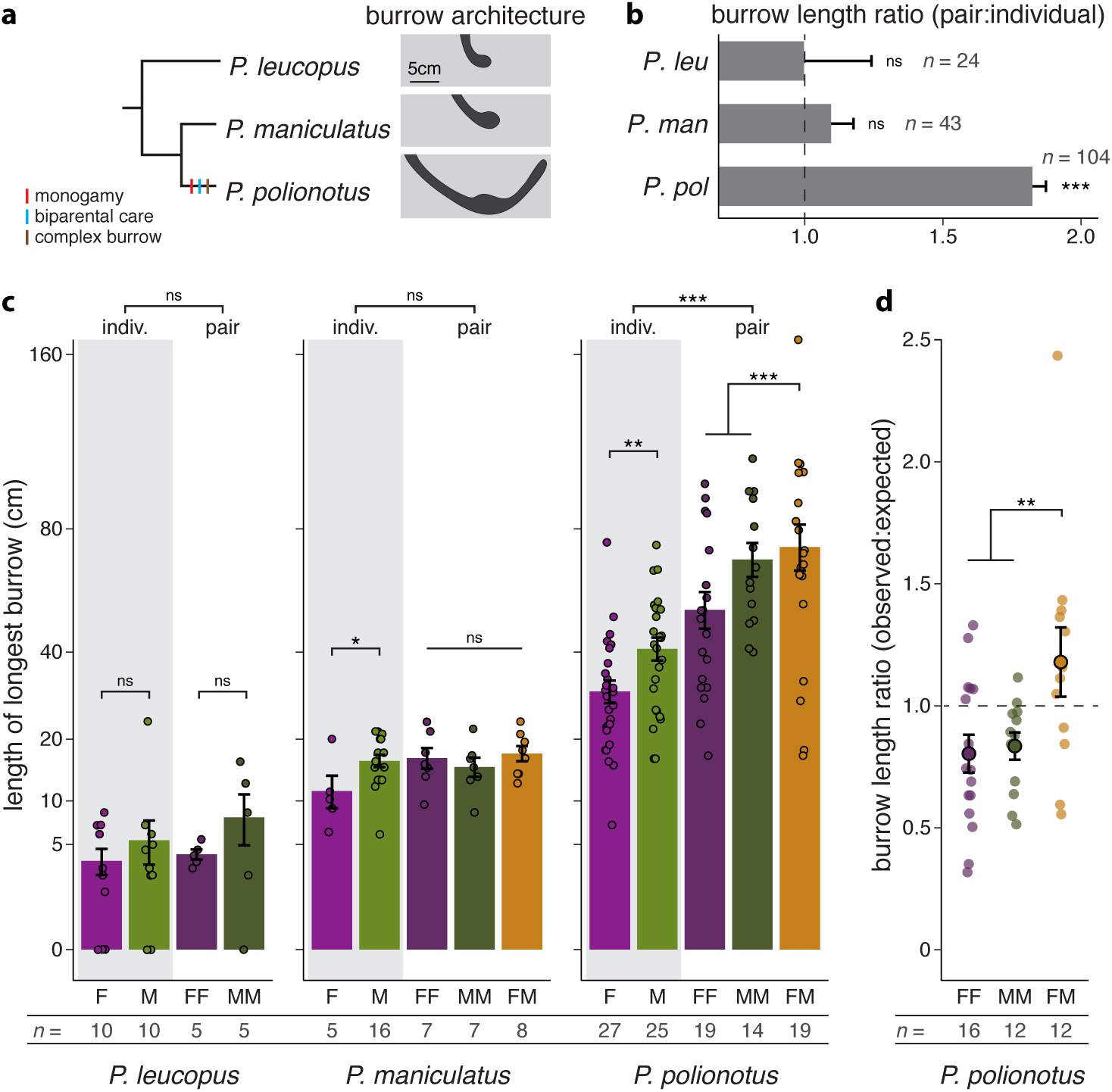
*P. polionotus* pairs cooperatively construct long burrows. **a**, Phylogenetic relationships, presence/absence of derived traits, and characteristic burrow architectures of three *Peromyscus* species. **b**, Ratio of pair to individual burrow lengths in three species. A ratio >1 indicates that two mice together dig a longer burrow than one mouse alone. All pair-types (FF, MM, FM) and both sexes (F, M) are included in ratio calculations (*P. leu*: *n* = 9 pairs, 15 individuals; *P. man*: *n* = 22 pairs, 21 individuals; *P. pol*: *n* = 52 pairs, 52 individuals). **c**, Length of longest burrow per trial dug by individuals (F = purple, M = green) and pairs of mice (FF = dark purple, MM = dark green, FM = gold) in three species. Data are plotte d on a square-root scale. Points represent the mean of up to 7 trials per individual or unique pair (mean 2.5 trials). **d**, Ratio of observed to expected burrow lengths in *P. polionotus* pairs, given the known output of individuals comprising the pair. Samples sizes are provided. \**P* < 0.05, \*\**P* < 0.01, \*\*\**P* < 0.001. Error bars represent s.e.m.

Because sex – of both the individual and one’s social partner – can be an important source of variation in behaviour, we next tested for sex differences in burrow length, both between individual males and females and between same- and opposite-sex pairs. In *P. leucopus*, we did not detect significant burrow length differences between sexes or pair-types (Fig. 1c, LMM, sex: *P* = 0.418, pair-type: *P* = 0.467). By contrast, in both *P. maniculatus* and *P. polionotus*, individual males dug longer burrows than individual females (Fig. 1c, LMMs, *P. man*: *P* = 0.027; *P. pol*: *P* = 0.001); in *P. maniculatus*, males dug burrows that were on average 42% longer, and in *P. polionotus*, males dug burrows that were 36% longer than females. However, only in *P. polionotus* did we observe variation in burrow length between pair-types, with opposite-sex pairs producing significantly longer burrows, on average 14 cm (or 28%) longer, than same-sex pairs (Fig. 1c, LMM, *P* < 0.001).

To determine if sex differences in burrowing behaviour can explain the observed variation among *P. polionotus* pair-types, we next tested whether burrows dug by different pair-types were shorter or longer than expected, given the length of the individual burrows dug by animals comprising the pair. For each *P. polionotus* pair-type, we calculated the observed:expected burrow length ratio, in which the observed value is the mean burrow length dug by a given pair, and the expected value is the sum of the mean burrow lengths produced individually by each member of the pair. We found that opposite-sex pairs dug longer burrows than expected compared to same-sex pairs (Fig. 1d, LMM, *P* = 0.005). The observed:expected ratio was significantly less than 1 for same-sex pairs, but not significantly different from 1 for opposite-sex pairs (Fig. 1d, *t*-tests, FF: *P* = 0.012; MM: *P* = 0.006; FM: *P* = 0.234), suggesting that *per capita* output declines with a same-sex partner but is unchanged with an opposite-sex partner. These data suggest that *P. polionotus* mice dig differently if they burrow with a same- or opposite-sex partner, raising the question if mice invest more when constructing a burrow intended for reproduction.

### Effect of reproduction on burrow length in females

Early studies of burrowing in *P. polionotus* hypothesized that reproduction may induce increased burrow construction in females^11^. Accordingly, we tested if females increase their burrow output in response to (or in anticipation of) a change in reproductive status, and, if so, if this could explain the observation that opposite-sex pairs of *P. polionotus* construct longer burrows. We used a repeated measures design to compare the burrows dug by individual females before and after being co-housed with a conspecific male (Fig. 2a). We tested for an effect of reproduction on burrow length by generating LMMs controlling for female age and length of the cohabitation period. Importantly, females with longer virgin burrows were no more likely to become pregnant than those with shorter virgin burrows, discounting a relationship between burrow construction and fertility (binary logistic regression, *P* = 0.833).

**Figure 2.**
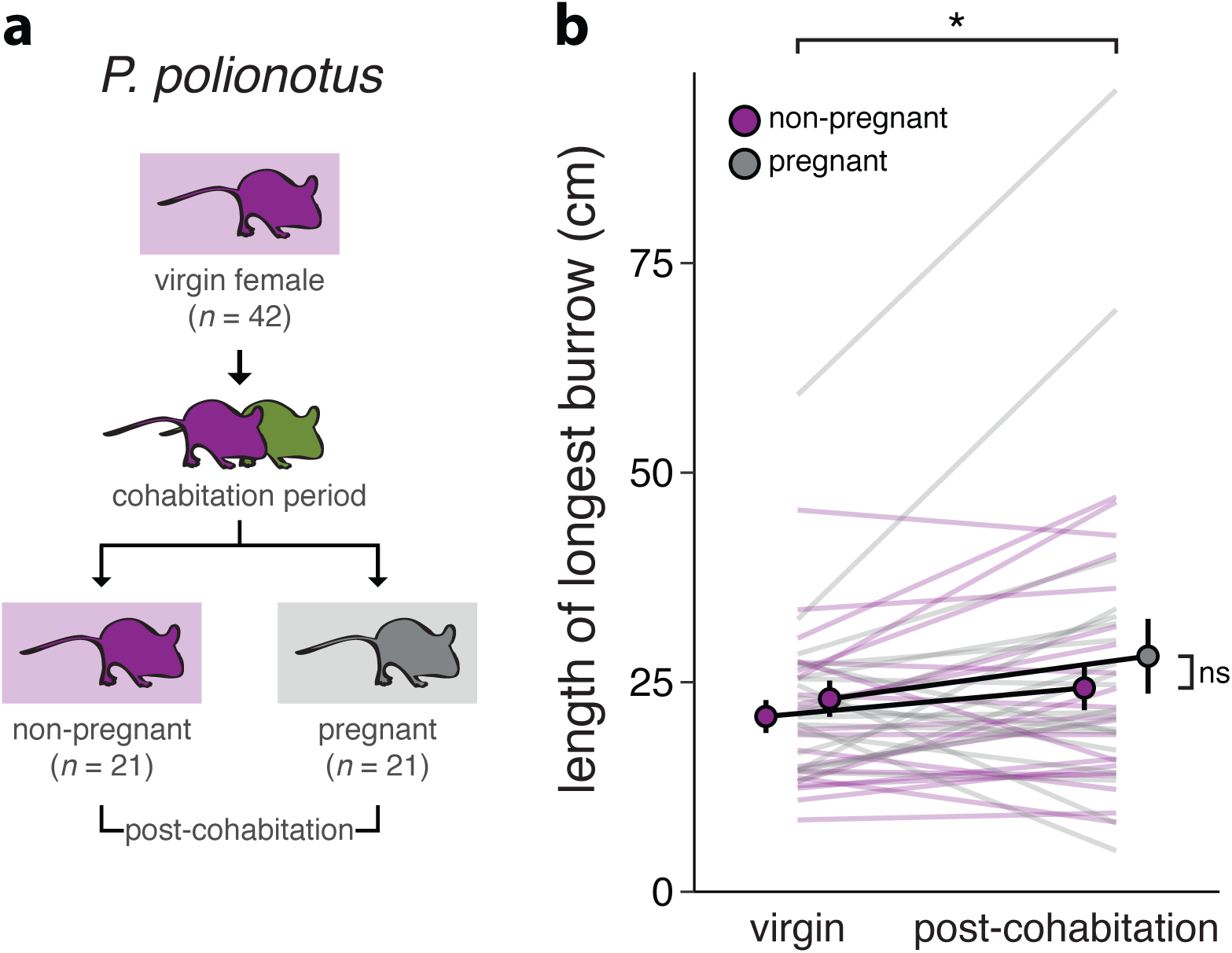
*P. polionotus* females dig longer burrows after male cohabitation. **a**, Experimental design schematic. Individual females were first tested as virgins to determine baseline burrowing output (top). After a period of cohabitation with a male (middle), during which half the mice became pregnant, females were individually tested again (bottom). **b**, Length of longest burrow dug by females before (left) and after (right) male cohabitation. Each line represents the mean of 2 trials per individual, per timepoint. \**P* < 0.05. Error bars represent s.e.m.

We found that, in *P. polionotus*, cohabitation with a male, but not pregnancy, impacted female burrow length. Regardless of pregnancy status, 26/42 (62%) of *P. polionotus* females produced slightly longer burrows after cohabitation with a male, with a median increase of 11% over their virgin trials (26.2 ± 2.6 cm vs. 22.0 ± 1.4 cm) (Fig. 2b, LMM, *P* = 0.023), but we found no difference in burrow length between pregnant and non-pregnant females (LMM, *P* = 0.320). We next asked if this 11% increase in burrow output by *P. polionotus* females exposed to males could explain the longer-than-expected opposite-sex burrows observed (see Fig. 1c). However, even after accounting for this increase, we still find a significantly greater observed:expected burrow length ratio for opposite-sex than same-sex pairs (LMM, *P* = 0.012). Thus, while cohabitation with a male can have modest effects on the burrowing behaviour of females, it does not explain the longer burrows produced by opposite-sex pairs of *P. polionotus*.

### Novel assay tracks individual behaviour

To uncover the behavioural mechanisms by which pairs of *P. polionotus* produce burrows of different lengths, we designed a narrow and transparent sand-filled enclosure in which we could directly observe and measure burrowing behaviour (Fig. 3a). With this assay, we first quantified individual digging behaviours and measured real-time progress in burrow construction (see Methods). Using a repeated measures design, we then tested if *P. polionotus* individuals alter their burrowing behaviour when digging with a same-versus opposite-sex partner (Fig. 3b). Specifically, we asked if the longer burrows produced by opposite-sex pairs result from increased individual effort (i.e., increased digging duration) with an opposite-sex partner.

**Figure 3.**
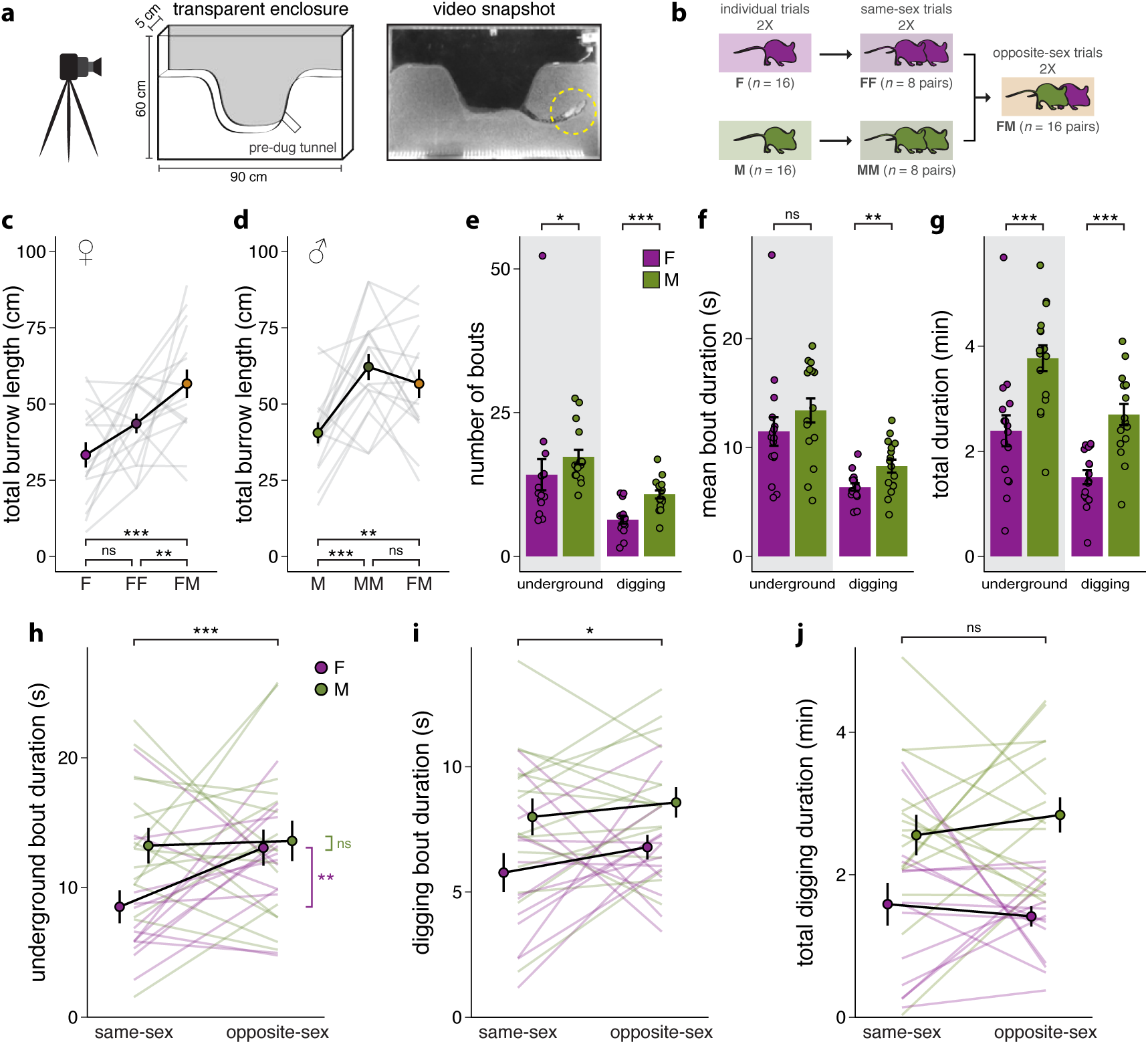
*P. polionotus* mice do not spend more time digging in opposite-sex trials. **a**, Schematic of the behavioural apparatus and snapshot of two mice digging simultaneously. **b**, Experimental design. Baseline burrow output was first quantified in virgins (left), then mice were tested in random order with both a same-sex (middle) and opposite-sex (right) burrowing partner. **c-d**, Total burrow length produced in three social contexts: individual, same-sex, and opposite-sex trials. Each line represents the mean of 2 trials per mouse, per context for females (**c**) and males (**d**). **e-g**, Underground and digging behaviour, pooled across same- and opposite-sex trials for individual females and males. Points represent the mean of 8 observations per individual. Benjamini-Hochberg adjusted *P* values are reported. **e**, Number of underground and digging bouts, per observation. **f**, Mean duration of underground and digging bouts, per observation. **g**, Total time spent underground and total time spent digging, per observation. **h-j**, Comparison of individual burrowing behaviour in same- and opposite-sex trials. Each line represents the mean of 4 observations per individual, per trial-type. **h**, Mean underground bout duration. **i**, Mean digging bout duration. **j**, Total digging duration. \**P* < 0.05, \*\**P* < 0.01, \*\*\**P* < 0.001. Error bars represent s.e.m.

To first confirm that burrowing behaviour in this new narrow enclosure is similar to that observed in the large enclosures, we measured the total burrow length dug by individuals and pairs of mice and tested for differences controlling for trial number and mouse identity. For females, burrowing with an opposite-sex, but not same-sex, partner significantly increased total burrow length (Fig. 3c; Tukey’s test, F vs. FF: *P* = 0.219; F vs. FM: *P* < 0.001). However, for males, burrowing with a partner of either sex significantly increased total burrow length (Fig. 3d; Tukey’s test, M vs. MM: *P* < 0.001; M vs. FM: *P* = 0.003). These results suggest that while males build similar burrows with a partner of either sex, females build longer burrows only with an opposite-sex partner, consistent with the pattern observed in the large enclosures (see Fig. 1c).

### Sex differences in burrowing behaviour

To identify differences in mouse behaviour that may contribute to sex differences in burrow length, we quantified individual-level behaviours using data from both same- and opposite-sex trials. Controlling for trial-type, we found that males had more underground bouts (i.e., entered the burrow more often) and more digging bouts (i.e., periods of continuous digging) than females (Fig. 3e, LMMs, underground: *P* = 0.021; digging: *P* < 0.001). While the duration of underground bouts did not differ between sexes, males had significantly longer digging bouts than females (Fig. 3f, LMMs, underground: *P* = 0.053; digging: *P* = 0.002). Overall, males spent more time underground and more time digging than females (Fig. 3g, LMMs, underground: *P* < 0.001; digging: *P* < 0.001).

To determine if individual contributions to shared burrows differ between same- and opposite-sex pairs, we calculated the division of labour (i.e., proportion of total digging time performed by each individual in the pair). Whereas the total time spent digging was split evenly between each member of a same-sex pair, the division of labour was significantly skewed in opposite-sex pairs (Fig. S1a, LMM, *P* < 0.001). On average, females performed 32% of the total digging time, while males contributed the remaining 68%.

Because males spend more time digging than females, we may expect (under a simple additive model) that two males would spend the most time digging and produce the longest burrows. However, surprisingly, opposite-sex pairs spend just as much time digging (Fig. S1b) and produce burrows that are just as long (Fig. S1c) as male-male pairs. These data from paired trials are compatible with the possibility that individual burrowing behaviour may differ with social context.

### Effect of social context on burrowing behaviour

To determine if social context affects individual-level behaviour, we compared burrowing behaviour between same- and opposite-sex trials using a repeated measures design (see Fig. 3b). We found that, on average, females had 53% longer underground bouts in opposite-sex trials, while males remained unaffected (Fig. 3h, least-squares means, females: *P* = 0.002, males: *P* = 0.999). Both sexes showed 10% longer digging bouts in opposite-sex trials (Fig. 3i, LMM, *P* = 0.039). Despite these differences, we nonetheless did not detect any differences in either the number of digging bouts (LMM, *P* = 0.112) or, importantly, the total digging duration between same- and opposite-sex trials for either sex (Fig. 3j, LMM, *P* = 0.340). Together, these results suggest that, although some behaviours differ slightly depending on social context, there is no detectable difference in total time spent digging by individuals in same-versus opposite-sex pairs. Thus, mechanisms other than increased individual effort (i.e., digging duration) are important for determining pair burrow length.

### Differences in social interactions between pairs-types

We next tested for differences in social behaviour that might indicate differences in social interactions (i.e., social cohesion). We found that opposite-sex pairs engaged in more affiliative behaviours (e.g., allogrooming, huddling) than same-sex pairs (Fig. 4a, LMM, *P* = 0.004). Furthermore, we observed agonistic interactions (e.g., boxing, biting, fleeing) in 44% of male-male trials, but only 28% of opposite-sex trials, while no agonistic interactions were observed in female-female trials (Fig. 4b, Fisher’s exact test, MM vs. FF: *P* = 0.0073; FM vs. FF: *P* = 0.041). These results raise the possibility that differences in social interactions among pair-types may reflect differences in the propensity to burrow cooperatively.

**Figure 4.**
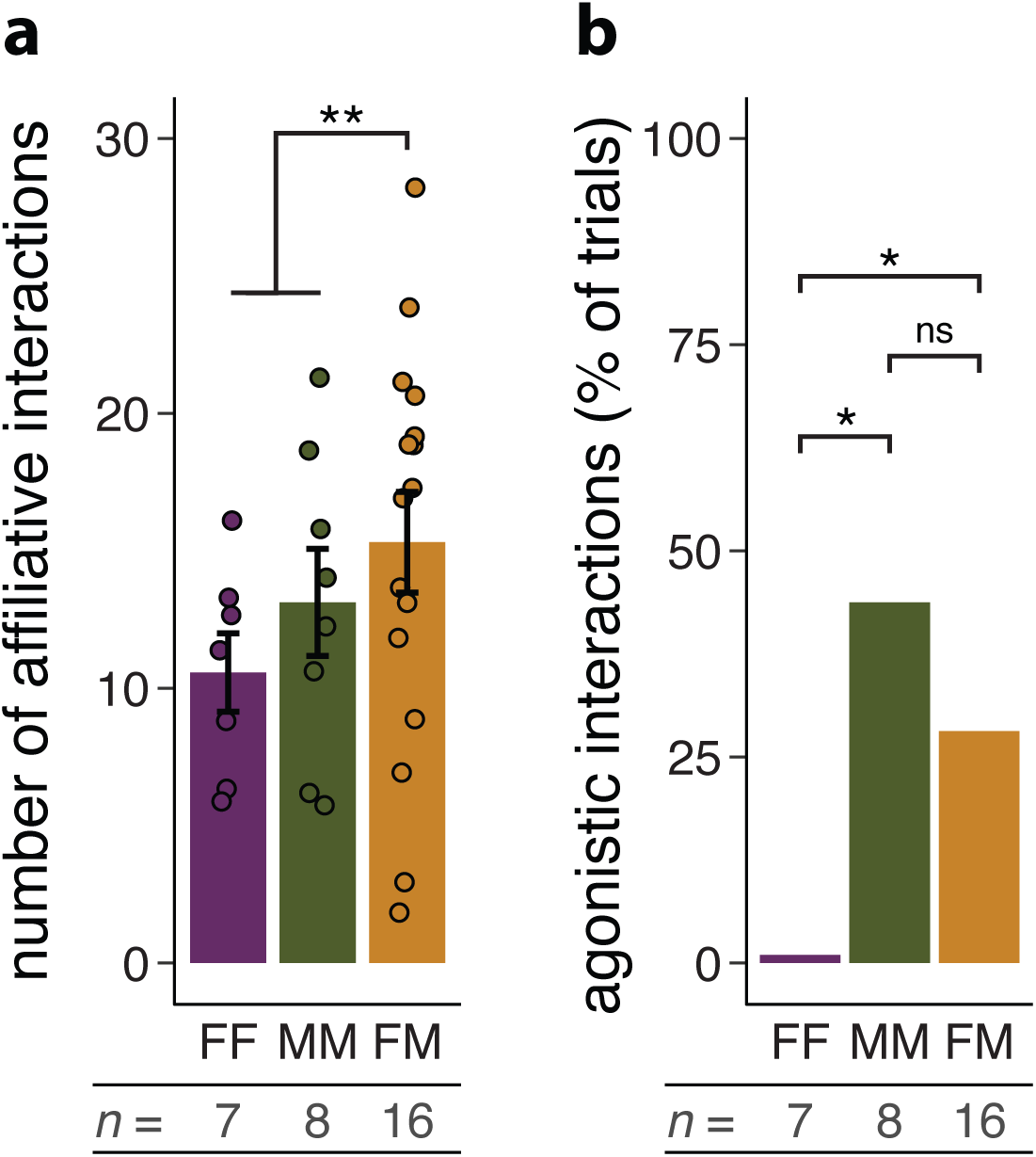
Opposite-sex pairs have more affiliative and fewer agonistic interactions. **a**, Total number of affiliative interactions observed per unique pair. Each point represents the sum of 4 observations per pair. **b**, Percentage of trials in which agonistic behaviour was observed between partners. Sample sizes are provided \**P* < 0.05, \*\**P* < 0.01. Error bars represent s.e.m.

### Differences in digging efficiency between pairs-types

To understand how opposite-sex pairs generated longer burrows despite no significant increase in digging duration, we tested for differences in efficiency among pair-types. To calculate digging efficiency (i.e., burrow extension rate), we divided the change in burrow length over the course of each 10-minute observation period by the total digging duration in that period (see Methods). We distinguished between independent digging (i.e., mice working independently, either temporally or spatially) and simultaneous digging (i.e., both mice working at the same place at the same time) (Video S1). As expected, the duration of both independent digging and simultaneous digging were significant predictors of change in burrow length (Fig. 5a). We calculated partial correlation coefficients for both independent and simultaneous digging (Fig. 5a, independent digging: *r* = 0.255, *P* < 0.001; simultaneous digging: *r* = 0.356, *P* < 0.001). To control for the number of mice, independent and simultaneous digging were both expressed in man-hours or, more accurately, mouse-minutes (i.e., time spent digging multiplied by the number of mice digging). We found that an additional mouse-minute of independent digging resulted in an additional 0.46 ± 0.13 cm of burrow length (Fig. 5b). However, an additional mouse-minute of simultaneous digging resulted in an additional 1.10 ± 0.21 cm of burrow length, indicating that concurrent, coordinated digging is a more efficient mode of burrow extension, even when controlling for total digging duration.

**Figure 5.**
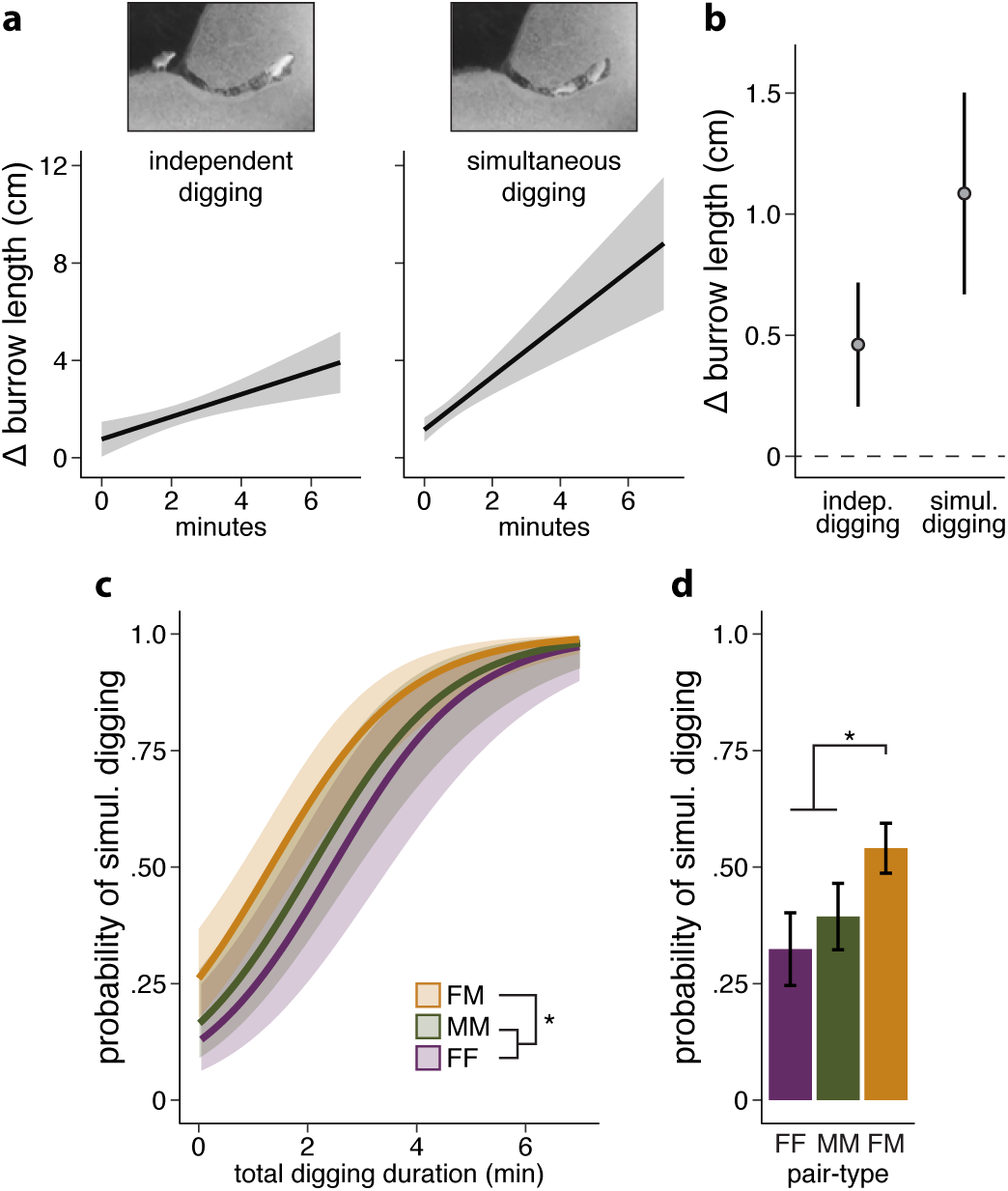
Opposite-sex pairs are more likely to engage in efficient simultaneous digging. **a**, Partial regression plots. Estimated relationship between independent digging duration (i.e., one mouse digging at a time) and change in burrow length, controlling for simultaneous digging time (left). Estimated relationship between simultaneous digging duration (i.e., both mice working concurrently) and change in burrow length, controlling for independent digging time (right). **b**, Mean effect estimates for independent and simultaneous digging duration on change in burrow length. Error bars represent 95% CI. **c**, Probability of simultaneous digging for simulated focal mouse digging durations, given a fixed partner mouse digging duration. **d**, Probability of observing simultaneous digging during same- and opposite-sex trials. \**P* < 0.05. Error bars represent s.e.m.

We then tested if the probability of simultaneous digging differed among pair-types. We found that opposite-sex pairs were more likely to engage in simultaneous digging, even after controlling for time spent digging by both the focal mouse and its partner (Fig. 5c, binary logistic regression, *P* = 0.025). The probability of observing simultaneous digging was significantly higher for opposite-sex than for same-sex pairs: 54 ± 5% for opposite-sex pairs, but only 39 ± 7% for two males and 32 ± 8% for two females (Fig. 5d, binary logistic regression, *P* = 0.025). Together, our results suggest that longer opposite-sex burrows can be explained predominately by an increase in burrowing efficiency mediated by enhanced social cohesion rather than by an increase in individual burrowing effort (i.e., digging duration).

## Discussion

The burrow is often a primary site for social interactions in many rodent species^16^ and is thus a particularly relevant context in which to assess social behaviour. Here, we capitalized on variation among *Peromyscus* mice to explore how different social contexts (i.e., digging alone versus with a partner) may alter burrowing behaviour across species with divergent mating systems.

We first found that *P. polionotus* mice, when paired, dig cooperatively and produce burrows that are longer than those built by individual mice. This pattern is consistent with what is known about their mating system and burrowing behaviour in the wild. Specifically, *P. polionotus* is both socially and genetically monogamous^22^, shows biparental care of young^23^, and in the wild most often nests in a burrow with an opposite-sex partner^17-19^. For example, field studies have shown that in two-thirds of excavated burrows, multiple mice (usually a male and a female), rather than a single individual, are found in the burrow^19^. In addition, in the wild, where mice are free to dig with a partner, burrows are longer than those produced by individuals in the laboratory^13^. By contrast, we found that both *P. maniculatus* and *P. leucopus* do not dig longer burrows as pairs than as individuals. These two species are thought to be largely promiscuous^24,25^, primarily display uniparental (i.e., maternal) care^23-26^, and most often nest solitarily, according to both radiotelemetry^27-29^ and nest-box studies^30,31^. Thus, this difference in social ecology may explain why we observed cooperative burrow construction only in *P. polionotus*, a species that typically cohabitates with a monogamous partner in the wild.

In *P. polionotus*, not only are jointly constructed burrows longer than those produced by individuals, but burrow length also differs depending on the sex of the paired individuals: opposite-sex pairs build longer burrows than same-sex pairs. It is tempting to imagine that mice alter their behaviour based on whether they are, for example, paired with a potential reproductive partner. In fact, early studies of burrowing hypothesized that females may alter their digging behaviour in response to (or in anticipation of) a change in reproductive status^11^. However, we found only marginal increases in burrow output after male cohabitation and no significant differences between pregnant and non-pregnant *P. polionotus* mice. Moreover, we show that mice do not spend more time digging in opposite-sex versus same-sex pairs. Thus, our initial prediction – that individuals actively increase their digging duration in response to an opposite-sex digging partner – was not supported.

Instead, we found that elongated burrows emerge largely because of greater spatiotemporal overlap between more socially cohesive opposite-sex pairs. In particular, opposite-sex pairs are more likely to dig simultaneously in the same burrow – a more efficient mode of burrow elongation that produces overall longer burrows despite no change in total digging duration. This simultaneous digging behaviour is reminiscent of “chain digging” observed in other communal burrowing mammals, such as eusocial mole rats^32^ and group-living degus^33^, and suggests that distantly related rodent species may have converged upon a common strategy for the efficient excavation of shared living space.

In other species that build collective structures, there is evidence that behaviour varies depending on social context. For example, in most ants and termites, nest volume correlates strongly with colony size^34^, and such adaptive scaling is mediated by a negative-feedback process in which insects adjust their digging rate according to traffic flow or the density of workers in the nest^35^. In *P. polionotus*, however, we find that while burrow size varies with pair-type, there is no direct change in digging behaviour (i.e., individual digging duration). This provides additional and compelling evidence that individual burrowing behaviour in *P. polionotus* is robust and largely innate, or at least invariant to social partnering. Thus, our study nicely highlights the importance of both designing novel behavioural assays to directly measure natural behaviour as well as dissecting the individual-level behaviours that collectively produce variation in social interactions and, in this case, ultimately communal structures.

Ernst Mayr^1^ may have predicted that, relative to the arguably less social species *P. maniculatus* and *P. leucopus*, monogamous *P. polionotus* would be more likely to have evolved an “open” genetic program to more easily adapt its behaviour to varying social environments. While this prediction may be true for other behaviours, burrowing in *P. polionotus* instead seems to conform nicely to what Mayr described as a fixed behaviour – that is, having a “closed” genetic program. Because individual burrowing behaviour appears largely impervious to perturbation (in this case, social context), it is a particularly promising trait for future genetic and neurobiological dissection to better understand the proximate mechanisms responsible for the evolution of innate behaviour.

## Acknowledgements

We thank N. Shen Molesky for help with behavioural assays; Z. Ali for helpful discussions; E. Hager, C. Hu, C. Lewarch, and O. Meyerson for comments on the manuscript. This work was supported by a James Milles Pierce Fellowship, Robert A. Chapman Memorial Scholarship, and a Natural Science and Engineering Research Council of Canada Postgraduate Scholarship to N.L.B.; a Robert A. Chapman Memorial Scholarship and a Harvard Mind Brain Behavior Graduate Student Award to J.N.W.; a Harvard Mind Brain Behavior Graduate Student Award to W.T.; an HHMI International Student Research Fellowship, Herchel Smith Graduate Fellowship, and a Joan Brockman Williamson Graduate Fellowship to F.B.; a Harvard College Program for Research in Science and Engineering Fellowship to A.K.; a Harvard College Program for Research in Science and Engineering Fellowship and a Herchel Smith Undergraduate Fellowship to R.A.G. H.E.H. is an Investigator of the Howard Hughes Medical Institute.

## Author Contributions

N.L.B., J.N.W., W.T., F.B. and H.E.H. designed the experiments. N.L.B., J.N.W., W.T., F.B., A.K. and R.A.G. performed the experiments. N.L.B. analysed the data. N.L.B. and H.E.H. wrote the manuscript with input from all authors.

## Materials & Correspondence

Correspondence and material requests should be addressed to H.E.H.

## Methods

### Animal husbandry

We performed experiments with *P. polionotus subgriseus* (PO stock), *P. maniculatus bairdii* (BW stock), and *P. leucopus* (LL stock) maintained as outbred colonies at Harvard University. Stocks were originally obtained from the *Peromyscus* Genetic Stock Center at the University of South Carolina. We maintained animals in ventilated cages measuring 7.75 × 12 × 6.5” (Allentown Inc., Allentown, NJ), which were furnished with 1/4″ Bed-o’Cobs bedding (The Andersons, Maumee, OH), a 2” square cotton nestlet (Ancare, Bellmore, NY), and a red polycarbonate hut (Bio-Serv, Flemington, NJ). We provided food and water *ad libitum*. Upon weaning at 23 days, mice were co-housed in same-sex groups of at most five conspecifics and fed irradiated Prolab Isopro RMH 3000 (LabDiet, St. Louis, MO). We fed all paired adults irradiated breeder chow: PicoLab Mouse Diet 20 (LabDiet, St. Louis, MO). We maintained mice on a 16h light:8h dark cycle at 22°C. The Institutional Animal Care and Use Committee at Harvard University approved all protocols.

### Behavioural assays

#### Large enclosures

We measured burrow architecture as described previously^13,36^. Briefly, we introduced individuals or pairs of mice into large PVC boxes (1.2 × 1.5 × 1.1m) filled with 700kg hydrated, hard-packed premium play sand (Quickrete, Atlanta, GA). We provided food and water *ad libitum* along with a cotton nestlet. We removed mice from the enclosures after one (pregnancy trials) or two (standard trials) overnight active periods. Then, we made casts of the resultant burrows using polyurethane filling foam (Hilti, Schaan, Liechtenstein). Next, a researcher blind to trial identity hand-measured burrow casts. After each trial, we removed all food, feces, nesting material, and disturbed substrate from the enclosure, and then rinsed the enclosure walls with water and turned over the sand to minimize residual odours.

#### Narrow transparent enclosures

We also assayed the burrowing behaviour of mice directly using an acrylic chamber (5cm × 0.9m × 0.6m) with a transparent Plexiglass face (see Fig. 3a). Using a pre-cut mould, we sculpted hydrated sand into two symmetrical 45° hills and excavated an 8cm pre-dug tunnel from one randomly selected hill to encourage burrowing in a consistent location. We outfitted the apparatus with an infrared illuminator frame that enabled video recording in the dark. We then introduced individuals or pairs of mice into the enclosures at the start of the dark cycle and removed animals the next morning, recording 8 hours of continuous video during the dark cycle. We provided food and water *ad libitum*. Photographs of the resultant burrows were taken at the end of the trial; details of photo and video analyses are provided below. Following each trial, we removed all food, feces, and disturbed substrate and rinsed the enclosure to minimize residual odours.

### Experimental design

#### Individual and pair burrowing differences across species

We first measured the burrows of the three *Peromyscus* species in individual and pair trials in the large enclosures. To control for past experience, we varied the order in which mice were tested in individual, same-sex, or opposite-sex trials. On average, each individual and each unique pair were tested twice. Mice were released into the enclosures at the start of the dark cycle and retrieved after two full overnight periods (∼36 hours).

#### The effect of pregnancy on female burrowing

To test whether pregnancy alters burrowing performance, we compared the burrows dug by females before and after being co-housed with a male. First, we tested the burrow output of virgin females (*n* = 42 *P. polionotus, n* = 36 *P. maniculatus*) in the large enclosures. Each female was assayed twice with 2 days rest between trials. We then transferred each female to a new cage with a conspecific male. Approximately half of all paired females subsequently became pregnant (21/42 *P. polionotus*, 19/36 *P. maniculatus*). Co-housed females were then tested again twice, as individuals, with 2 days rest between trials. Females were returned to their home cage after each trial and monitored daily for parturition for up to 23 days. To minimize stress to pregnant females, all mice in this experiment were tested for one overnight period (∼18 hours) in the large enclosures.

#### Social modulation of burrowing behaviour in P. polionotus

To test whether *P. polionotus* mice modulate their behaviour in response to a digging partner, we used a repeated measures design. We video-recorded all trials in the narrow transparent enclosures (see Fig. 3a). To distinguish individuals, we shaved a patch of hair from both flanks of one randomly-selected member of the pair. Shaving was completed at least a week before the first pairing, and markings did not grow in over the course of the experiment. To quantify baseline individual burrow output, we first assayed each individual twice (*n* = 16 virgin females, *n* = 16 virgin males). Next, we transferred each individual to a new cage with an unrelated and unfamiliar partner of the same or opposite sex. Pairs were given 2 nights to acclimate before being tested together in the small transparent enclosures. We recorded 8 hours of video for each overnight trial. Pairs were then returned to their home cage for 3 nights rest before their second trial. Mice were then re-partnered and the process was repeated. To control for previous experience, we randomly assigned the order in which mice burrowed with a same-sex or opposite-sex partner. Females were monitored daily for parturition for up to 23 days after the end of the experiment. No females were pregnant at any point during the experiment.

### Behavioural Analyses

#### Photo analysis

We measured the length of burrows produced by individuals or pairs in the narrow enclosure using Fiji image processing software^37^. Each photo was measured separately by two researchers blind to trial identity. Burrow length measurements were highly correlated between researchers (Pearson correlation, *r* = 0.97, *P* < 0.001). We therefore used the mean of two measurements for all statistical analyses. To calculate digging efficiency, we also took video snapshots at the beginning and end of each 10-minute observation timepoint. From these snapshots, we measured burrow length as described above.

#### Video analysis

In 75% of trials, pairs had fully completed at least one burrow 3 hours after lights out. Thus, to target the active burrow construction phase, we selected two 10-minute timepoints, spaced 1 hour apart, in this 3-hour window for observation: 0:30–0:40 and 1:30–1:40 hours. In 5 of the 62 videos, we did not observe digging during the pre-selected timepoints. For these videos, we shifted our two timepoints, still spaced one hour apart, to later in the night. We accounted for this shift in all statistical models.

We quantified the behaviour of each mouse in a pair using The Observer XT Version 12.0 (Noldus, Leesburg, VA). Each 10-minute timepoint was randomly assigned to a researcher blind to trial identity for behavioural scoring according to the following scheme: for each individual, we scored all burrow entries or exits, as well as extending (i.e., forelimb digging at the growing end of the burrow), widening (i.e., forelimb digging at any other location in the burrow), and hind-kicking (i.e., vigorous, coordinated hindlimb movements that expel loosened sand from the burrow) (Fig. S3). For each pair, we also scored affiliative and agonistic behaviours, which could occur either inside or outside the burrow. We defined affiliative behaviours as beneficial social interactions (i.e., allogrooming, huddling). We defined agonistic behaviours as aggressive interactions (i.e., boxing, parrying, biting) or submissive behaviours (i.e., freezing, fleeing). Altogether, each mouse (*n* = 32) received 80 minutes of direct observation across the entire study – 10 minutes per timepoint, 2 timepoints per trial, 2 trials per trial-type, 2 trial-types (same-sex and opposite-sex). We excluded observations from one same-sex pair in which the markings of the two females were indistinguishable and ID could not be confidently assigned.

### Statistics

All statistical tests were performed in R Version 3.2.3 (R Core Team, 2015).

#### Individual and pair burrowing differences across species

To normalize the data, we first square-root transformed burrow lengths. To test if pairs of mice dig longer burrows than individuals, we used a linear mixed-effects model (LMM) with social context (i.e., individual or pair trial) and sex as fixed effects and mouse ID as a random effect. We excluded trials in which individuals or pairs did not burrow. For each species, we divided the length of the average burrow dug by two mice by the length of the average burrow dug by one mouse to determine the pair:individual burrow length ratio (*n* = 9 pairs, 15 singles (*P. leu*); 22 pairs, 21 singles (*P. man*); 52 pairs, 52 singles (*P. pol*) (Fig. 1b). We approximated the variance of each ratio by Taylor series expansion. To further evaluate sex and pair-type differences within species, we used LMMs with either sex or pair-type as a fixed effect and mouse ID as a random effect (Fig. 1c). Differences in observed:expected burrow length ratio between same-sex and opposite-sex pairs were evaluated by LMM, with mouse ID and partner ID as random effects (Fig. 1d).

#### The effect of pregnancy on female burrowing

Data were log-transformed to improve normality. To test if individual females dig longer burrows after being co-housed with a male, we used LMMs with timepoint (i.e., before or after male cohabitation), status (i.e., non-pregnant or pregnant), and their interaction as fixed effects and mouse ID, age, and length of cohabitation period as random effects. The non-significant interaction term was subsequently removed from the *P. polionotus* model. For *P. maniculatus*, we used a least-squares means post-hoc test to further evaluate the timepoint by status interaction. For both species, we used binary logistic regression to confirm that females with longer virgin burrows were not more likely to become pregnant.

#### Social modulation of burrowing behaviour in P. polionotus

When necessary, data were transformed to improve normality. To test the effect of social context on burrow length, we used LMMs with trial-type (i.e., individual, same-sex, or opposite-sex) as a fixed effect. We included mouse ID and trial number as random effects to control for repeated measures and whether mice first burrowed with a same-sex or opposite-sex partner.

#### Sex differences

Because we found no sex or trial-type differences in the proportion of burrow extending versus burrow widening behaviour (LMM, sex: *P* = 0.490, trial-type: *P* = 0.384), we collapsed these two categories into one general “digging” designation. To assess overall sex differences in digging behaviour, we used LMMs with mouse sex and trial-type (i.e., same- or opposite-sex) as fixed effects and mouse ID, observer ID, trial number, and timepoint as random effects. To test explicitly for social modulation of digging behaviour, we used LMMs with mouse sex, trial-type, and their interaction as fixed effects and mouse ID, observer ID, trial number, and timepoint as random effects. For the underground bout model, we used a least-squares means post-hoc test to further evaluate the sex by trial-type interaction.

#### Pair-type differences

To calculate division of labour, we divided the total digging duration for each mouse by the total digging duration for the pair. Individuals in same-sex pairs were distinguished by their markings (i.e., shaved or non-shaved) and individuals in opposite-sex pairs were distinguished by sex. We used an LMM to test for a difference in division of labour between same- and opposite-sex pairs, including mouse ID, observer ID, trial number, and timepoint as random effects (Fig. S1a). We used LMMs and Tukey’s post-hoc comparisons to evaluate differences in total digging duration and total burrow length among pair-types (Fig. S1b, c). For total digging duration, we included pair ID, observer ID, trial number, and timepoint as random effects in the LMM. For total burrow length, we included pair ID and trial number as random effects.

#### Differences in social behaviour

To quantify differences in social interaction among pair-types, we summed the number of affiliative encounters per pair across all observations. We then used an LMM with pair-type as a fixed effect and pair ID, observer ID, trial number, and timepoint as random effects. Since Tukey’s test revealed no significant difference between FF and MM (*P* = 0.726), we re-ran the above LMM with trial-type (i.e., same- or opposite-sex) as a fixed effect rather than pair-type (i.e., FF, MM, or FM). Agonistic interactions were much rarer. We therefore counted the number of trials with and without any observed agonistic interactions and used pairwise Fisher’s exact tests to evaluate differences among pair-types.

#### Modelling burrowing efficiency

We modelled the relationship between independent digging, simultaneous digging, and change in burrow length using multiple linear regression. We then used binary logistic regression to test if opposite-sex pairs were more likely than same-sex pairs to engage in simultaneous digging, controlling for the time spent digging by both the focal mouse and its partner.

---

**Figure S1.**
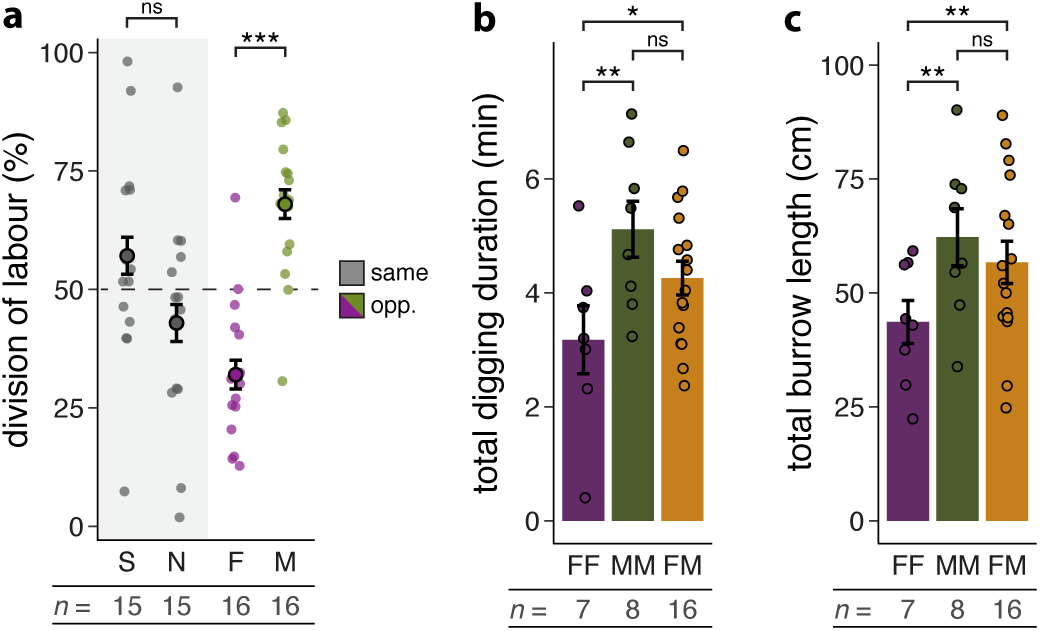
Pair-type differences in division of labour, total digging duration, and total burrow length. **a**, Division of labour in same- and opposite-sex pairs. On average, each mouse in a same-sex pair contributed equally to the total time spent digging (LMM, *P* = 0.096, *n* = 30), whereas the division of labour in opposite-sex pairs was significantly skewed (LMM, *P* < 0.001, *n* = 31). For same-sex pairs (grey), points on the left represent shaved (S) mice and points on the right represent non-shaved (N) mice. Points represent the mean of 4 observations per individual, per trial-type. **b**, Total time spent digging per observation by both mice in a pair. Opposite-sex pairs and male-male pairs spent more total time digging than female-female pairs (Tukey’s test, FM vs. FF: *P* = 0.039; MM vs. FF: *P* = 0.004, FM vs. MM: *P* = 0.406). Points represent the mean of 4 observations per pair. **c**, Total burrow length dug by three different pair-types. Opposite-sex pairs and male-male pairs built longer burrows than female-female pairs (Tukey’s test, FM vs. FF: *P* = 0.003; MM vs. FF: *P* = 0.006; FM vs. MM: *P* = 0.491). Points represent the mean of 2 trials per pair. Sample sizes are provided. \**P* < 0.05, \*\**P* < 0.01, \*\*\**P* < 0.001. Error bars represent s.e.m.

**Figure S2.**
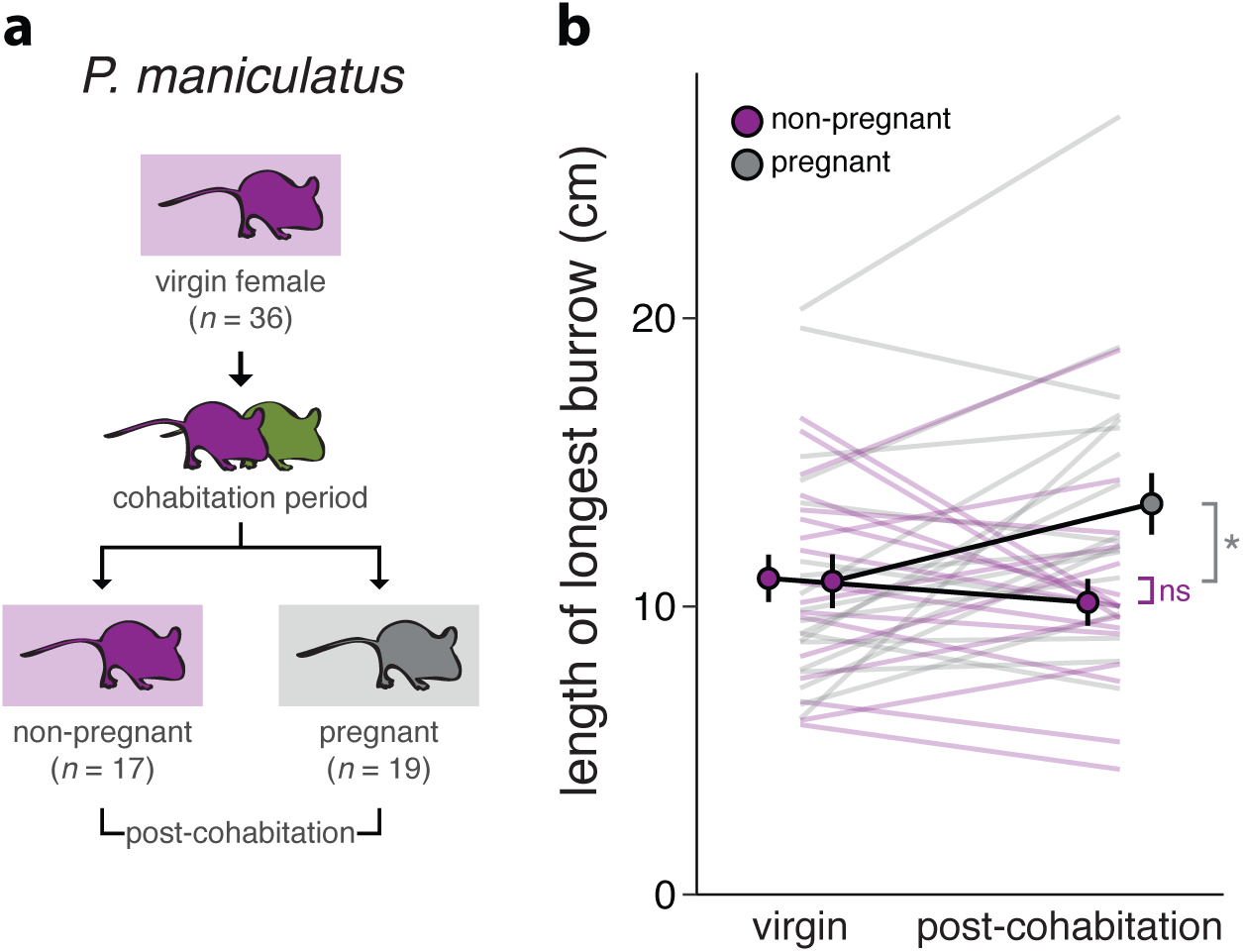
*P. maniculatus* females dig longer burrows when pregnant. **a**, Experimental design schematic. Individual females were first tested as virgins to determine baseline burrow output (top). After a period of cohabitation with a male (middle), during which 19/36 mice became pregnant, females were individually tested again (bottom). **b**, Length of longest burrow dug by females before (left) and after (right) male cohabitation. After male cohabitation, 79% (15/19) of pregnant females (grey) dug longer burrows, compared to only 35% (6/17) of non-pregnant females (purple). The median burrow length increase among pregnant females was 21% (least-squares means, *P* = 0.023, *n* = 19). By contrast, females that did not become pregnant after male cohabitation showed no change in burrow length relative to their virgin trials (least-squares means, *P* = 0.787, *n* = 17). Each line represents the mean of 2 trials per individual, per timepoint. \**P* < 0.05. Error bars represent s.e.m.

**Figure S3.**
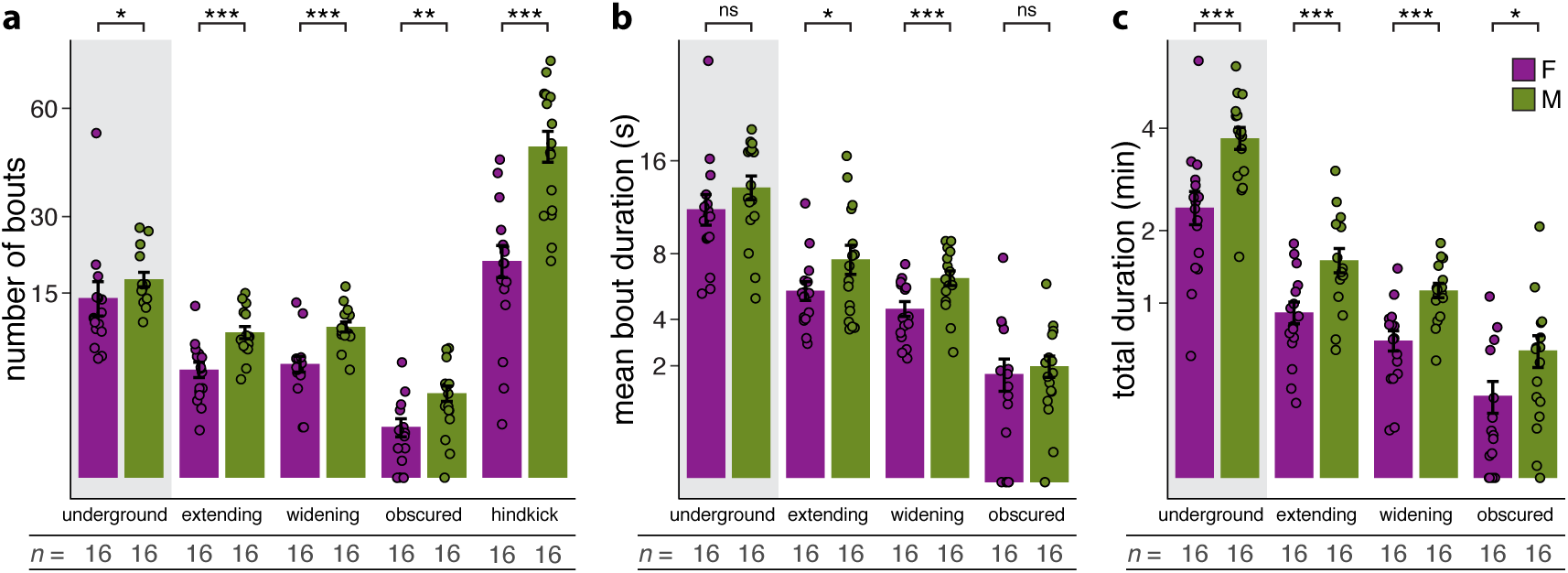
Sex differences in individual burrowing behaviours. **a-c**, Underground and digging behaviour, pooled across same- and opposite-sex trials for individual females and males. Behaviour labels are as follows: “underground” (i.e., mouse is in the burrow), “extending” (i.e., mouse is digging at the leading edge of the burrow), “widening” (i.e., mouse is expanding the interior of the burrow), “obscured” (i.e., mouse is underground but not visible), “hindkick” (i.e., expulsive kick that ejects loose sand from the burrow). Points represent the mean of 8 observations per individual. Benjamini-Hochberg adjusted *P* values are reported. **a**, Number of bouts, per observation. **b**, Mean bout duration, per observation. **c**, Total duration of behaviour, per observation. Sample sizes are provided. \**P* < 0.05, \*\**P* < 0.01, \*\*\**P* < 0.001. Error bars represent s.e.m.

